# ILK play a key role in partial bladder outler obstruction (PBOO) by regulation TLR4/NF-κB(p65) pathway

**DOI:** 10.1101/2021.04.26.441552

**Authors:** Yiduo Zhou, Yi Huang, Jie Gao, Le Shu, Sicong Zhang, Zhengsen Chen, Baixin Shen, Zhongqing Wei, Liucheng Ding

## Abstract

**Aim:** The purpose of this research was to discuss the effects and relative mechanisms of ILK in PBOO by vivo and vitro study.

**Materials and methods:** The SD rats were divided into Normal, Sham and Model groups. Collecting Bladder outlet tissue, observation pathology and fibrosis levels by H&E and Masson staining. Measuring cell apoptosis and cell viability by TUNEL and p-histone H3 staining, ILK protein were evaluated by WB and IHC assay in Bladder outlet tissue. Using TGF-β1 stimulating BSMC cell to make PBOO cell model. Measuring cell proliferation by CCK-8 assay; Relative gene and proteins expression were evaluated by immunofluorescence, WB and RT-qPCR assay.

**Results:** Compared with Normal group, bladder weight, collage fiber area, apoptosis cell number and cell viability were significantly difference with ILK protein significantly increasing in bladder outer tissues of Model group (P < 0.05, respectively). In vitro cell experiment, ILK overexpression had effects to stimulate cell proliferation via TLR4/NF-κB(p65) pathway; however, with ILK knockdown, the cell proliferation was significantly depressed via regulation TLR4/NF-κB(p65).

**Conclusion:** ILK play an important role in PBOO induced cell proliferation, ILK knockdown had effects to improve PBOO induced cell hyper-proliferation via depressing TLR4/NF-κB(p65) pathway.

## Introduction

Partial Bladder Outlet Obstruction (PBOO) is defined as morphological and functional abnormality concerning bladder due to obstruction of any part located between bladder neck’s outlet and external urethral orifice as a result of a variety of conditions like Benign Prostatic Hyperplasia (BPH) and urethral stricture [1]. Now the goal of conventional treatment is to relieve the urinary obstruction largely by virtue of drugs and surgery which can achieve the goal but fails to improve the Lower Urinary Tract Symptoms (LUTS) among many patients. Therefore, more and more researchers are shifting their attention from obstruction relief to handling of PBOO-induced bladder dysfunction. Researches indicate apart from obstruction, the pathological changes of bladder secondary to obstruction remains more closely related to LUTS. This accounts for the significance of understanding better the molecular mechanism behind the abnormal biological behaviors of bladder smooth muscle in terms of its morphology and function after occurrence of obstruction.

In accordance with findings of existing studies, obstruction-induced excessive smooth muscle cell proliferation of bladder is the primary reason why bladder function cannot be restored [3]. As revealed by previous studies, phenotypic transformation of smooth muscle cell arising from bladder impairment plays a critical part in the genesis and development of human diseases [4,5]. However, the molecular mechanism underlying the phenotypic transformation of Bladder Smooth Muscle Cell (BSMC) remains still unclear. A variety of signal molecules, including PDGFPDGF-BB, TGFTGFβ, ECM and such environmental factors as ischemia, anoxia, bladder pressure, and mechanical stretch, all play a significant role in regulating BSMC phenotypic transformation and proliferation [5].

As researches indicate, ILK-mediated signal pathway plays a vital part in ventricular outflow tract obstruction and stays as an important upstream regulating mechanism behind overloading hypertrophy [6]. By contrast, in non-myocardial cells, when ILK gets activated, a series of cascade reactions leading to protein kinases’ phosphorylation will be initialized. The protein kinases hereby include PKB, GSK3, GSK3β, MAPK, ERKs and mTOR, which all stay closely concerned with BSMC growth [7]. It can be reasonably assumed that ILK matters a lot in causing PBOO.

## Materials and Methods

### Materials

Male SD rats were purchased from Zhejiang Charles River Laboratories (CRL, SCXK (ZHE) 2019-0001). Other instruments and reagents are listed as follows: hematoxylin-eosin dye kit (KeyGEN BioTECH KGA224, Jiangsu, China), Masson dye kit (Yike Biological, Guangzhou, China), TUNEL detection kit (KeyGEN BioTECH KGA702, Jiangsu, China), mouse anti-TnT (Abcam ab8295, Britain), rabbit anti-phospho-HistoneH3 (Abcam ab32107, Britain), TRITC goat anti-mouse (Jackson ImmunoResearch 115-025-062, U.S), FITC goat anti-rabbit (Jackson ImmunoResearch 111-095-003, U.S), ILK (KeyGEN BioTECH KGYT2346-7, Jiangsu, China), TRIzol (Invitrogen 15596-026, U.S), cDNA first strand synthesis kit (TaKaRa RR036B, Japan), and One Step TB Green™ PrimeScript™ RT-PCR Kit II (SYBR Green) (TaKaRa RR086B, Japan).

#### PBOO rat model

The SD rats underwent 12h fasting before being allowed to acclimate for 1 week. Then, they were intraperitoneally anesthetized with 10% chloral hydrate (0.3ml/00g) first, then fixed at supine position to have inferior belly skin prepared. Vertical incision (1.5-2.0cm) was formed in the right middle of iodine-sterilized inferior belly. Lateral rectus blunt dissection was done to open the abdomen and expose the bladder. One1mm epidural catheter was placed right next to the bladder neck without undermining the latter, and bladder neck was threaded with 3-0 suture to have it ligated upon the support of the epidural catheter. Later, the catheter was withdrawn and incision was blocked layer by layer. After 2 weeks’ routine breeding, the rats were finally executed for sampling purpose.

### H&E staining

Dewaxing and hydration were conducted as conventionally. The sections were then immersed in xylene twice, 5min at each; and dehydrated with ethyl alcohol at gradients. At last, dyeing was finished as specified by HE kit manufacturer.

### Masson staining

After conventional slicing was done, samples were stained with hematoxylin for 5min and then rinsed with distilled water, differentiated with 0.1% HCl and washed with tap water for making it turn blue again. Acidic ponceau dying liquid was used to dye the samples for 5min in order to restore the red color. After that, samples were rinsed with distilled water, treated with 1% phosphomolybdic acid solution for 50s, spin dried, stained with green color for 5s, rinsed completely with tap water, and dried in an electric oven. Once being dried, the sections were added with neutral gum and covered with slides and then examined and photographed under microscope. Fibrosis rates of bladder tissues in different groups were analyzed using Image J software.

### TUNEL staining

After conventional slicing was done, each 90μL 1×PBS sample was added with 10μL 10×ProteinaseK as needed to allow for reaction at 37°C for 15min. The samples were treated with 3% H_2_O_2_-methanol solution for 15min, then immersed in PBS for 5min and rinsed thrice. Afterwards, each sample was dropwise added with 10μL TdT enzyme reaction liquid to undergo 1h reaction in a dark and humid environment at 37°C, then immersed in PBS for 5min and rinsed thrice. 100μL Streptavadin-HRP was added dropwise to promote reaction in a dark and humid place at 37°C for 30min. The solution was then immersed with PBS for 5min, rinsed thrice, colored with DAB, re-stained with hematoxylon. Finally, samples were counted under optical microscope.

### p-HistoneH3 staining

After being generated in the conventional way, each section was added with 2 drops of 3% H_2_O_2_-methanol solution and blocked at ambient temperature (or 15°C) for 10min, and then rinsed with PBS thrice. After that, 50μL ready-to-use goat serum was dropwise added for incubation at ambient temperature for 20min. 50μL diluted primary antibody was dropwise added for another incubation in a humidified box at 37°C for 2h. PBS rinsing was performed thrice before 50μL secondary antibodies including FITC (diluted at 1:200) and TRITC (diluted at 1:200) for incubation in a dark environment for 1h. PBS rinsing was performed thrice for a third time. Re-staining, sealing and microscopic examination were carried out in turn.

### Immunohistochemical (IHC) staining

After conventional slicing and antigen retrieval were done, each section was added with 2 drops of 3% 3%H_2_O_2_-methanoy solution for 10min blocking at ambient temperature (25°C). Then, PBS rinsing was performed for 3 times, 3min at each time. The section was blocked and then added dropwise with 50μL goat anti-rabbit/mouse polymer for incubation within a humidified box at ambient temperature for 20min. PBS rinsing was performed again for 3 times and min at each time. Developing, re-staining, dehydration, blocking and microscopic examination were carried out in turn. Finally, the sections were analyzed using Image J software.

### Western blotting (WB) assay

Tissues or cells were extracted to be decomposed by 1mL Lysis Buffet. Then, they were placed on a shaking table at 4°C for a mild vibration for 15min. The centrifugation operation lasted 15min at 14,000rpm and 4 °C. Supernatant was collected to have the protein be quantified (through BCA). 100g/L SAS-PAGE was performed, followed by transferring and blocking. Primary and secondary (1: 1000) antibodies were added respectively for incubation. Developing was achieved in the way of enhanced chemiluminescence. Labwork4.0 image analysis software finished scanning. The protein expression was expressed as the ratio of target protein absorbance to GAPDH absorbance.

### Extraction and separation of rats’ primary BSMC cells

Adult rats were anesthetized and detrusor on bladder was collected. Urothelium, lamina propria, serosa and adipose tissue were carefully removed, while the middle layer of smooth muscle tissue was reserved. The resulting primary tissues were cut into some 1mm3 pieces that were digested with pancreatin/EDTA at 37°C for 30min. The pieces were then further cut into paste and placed in 0.1% type I collagenase for digestion at 37 °C for 30min. The resulting cell suspension was filtered with 200-mesh strainer. Cells were collected from the filtrate through centrifugation, PBS rinsing was performed twice. DMEM/F12 medium containing 10% FBS was used to re-suspend the cells, and the medium was changed every 2-3 days.

### Cell immunofluorescence

Cell samples (cell smears or coverslips) were air dried and then immersed in 4% paraformaldehyde fixer for 30min or overnight to improve the cell permeability. PBS rinsing was conducted thrice and 3min at each time. Each section was added with 2 drops of 3% H2O2-methanol solution, blocked at room temperature (15-25°C) for 10min, and rinsed with PBS for three times. 100μL ready-to-use goat serum was added dropwise for incubation at ambient temperature for 20min. 50μL primary antibody (diluted at 1:100) was added for incubation in a humidified box for 2h and rinsing with PBS for three times. Then 50μL secondary antibody FITC (diluted at 1:100) was added for incubation in a dark place at 37°C for 1h. PBS rinsing was performed for three times. After re-staining and blocking were done, microscope was employed to examine the expression of protein in cells. Three high-expression areas were photographed as records.

### Real-time fluorescence quantitative PCR (RT-qPCR) assay

TRIzol was used to extract total DNA of tissues or cells, and Prime ScriptTM first strand cDNA synthesis kit to reversely transcript it to cDNA which later acted as template for amplification. The primer sequence is provided in Table 1. Real-time fluorescence quantitative PCR assay was conducted and mRNA’s relative expression was computed using 2^−ΔΔCT^ method.

**Table 1.**
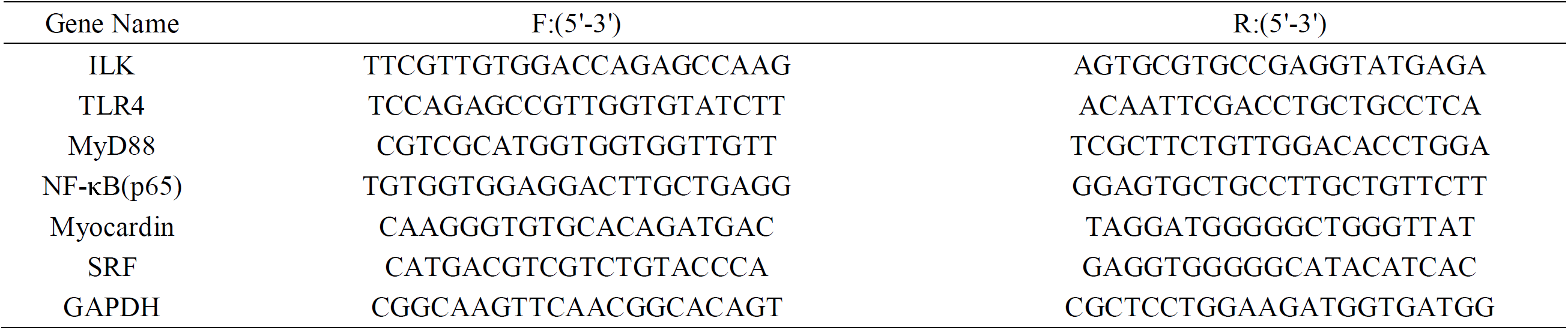
Primer sequence to RT-qPCR.

#### CCK-8 assay

Cells were digested, counted and formulated into 5×104/mL cell suspension. A 96-well cell culture plate was added with 100μL cell suspension in each well. The plate was placed into a cultivator containing 5% CO^2^ at 37◻ for 24h. The drug was diluted with complete medium to achieve the target concentration. Each well was filled with 100μL corresponding drug-containing medium. A negative control group was set up. The 96-well plate was then placed into a cultivator containing 5% CO^2^ at 37◻ for 24h. Each well was added with 10μL CCK-8 and further incubated for 3h before being mixed well on a shaking table for 10min. With λ=450 nm, the OD value of each well was read with a microplate reader in order to figure out the inhibition rate.

### Cell transfection

Primary BSMC cells were passed for two generations before being transfected. Both si-ILK and LV-ILK were designed and constructed by Nanjing KeyGEN BioTECH. The ILK-siRNA sequence is as shown in Table 2.

**Table 2.**
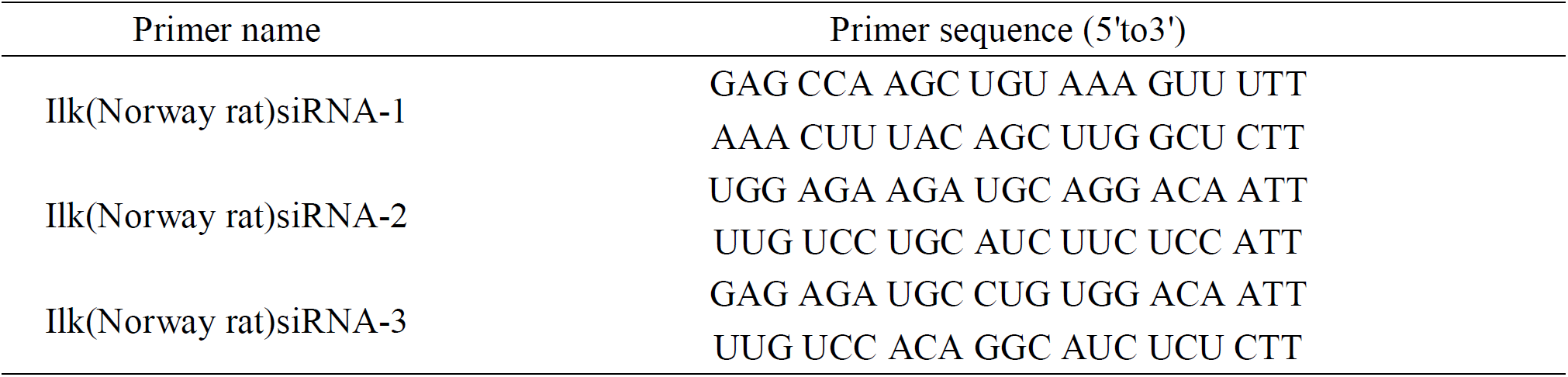
Primer sequence to siRNA.

### Statistical approach

Data statistics and analysis were performed using SPSS 22.0 software. All the measurement data were expressed as mean ± standard deviation (mean ± SD). Pairwise comparison adopted t test, while intergroup comparison was analyzed with one-way ANOVA. P<0.05 was considered as significant.

## Results

### Body weight, bladder weight and pathological changes of rats in different groups

As indicated by Fig. 1A, rats in different groups were insignificantly different in term of body weight. Sham group was not significantly different from the Normal group in bladder weight, but Model group witnessed a significant rise in bladder weight (P<0.05). according to HE staining results, nothing abnormal was detected in the detrusor cells of Normal group or Sham group, but not in the Model group as the last group presented detrusor cell fusion, less detrusor cells, more vacuolar degeneration, massive infiltration by inflammatory cell, accumulating extra-cellular connective tissue and more evident bladder fibrosis (Fig. 1B). Masson staining revealed smaller collagenous fiber area in Normal and Sham groups and significantly larger collagenous fiber area in Model group than in Normal group (P<0.001, Fig. 1C).

**Figure 1.**
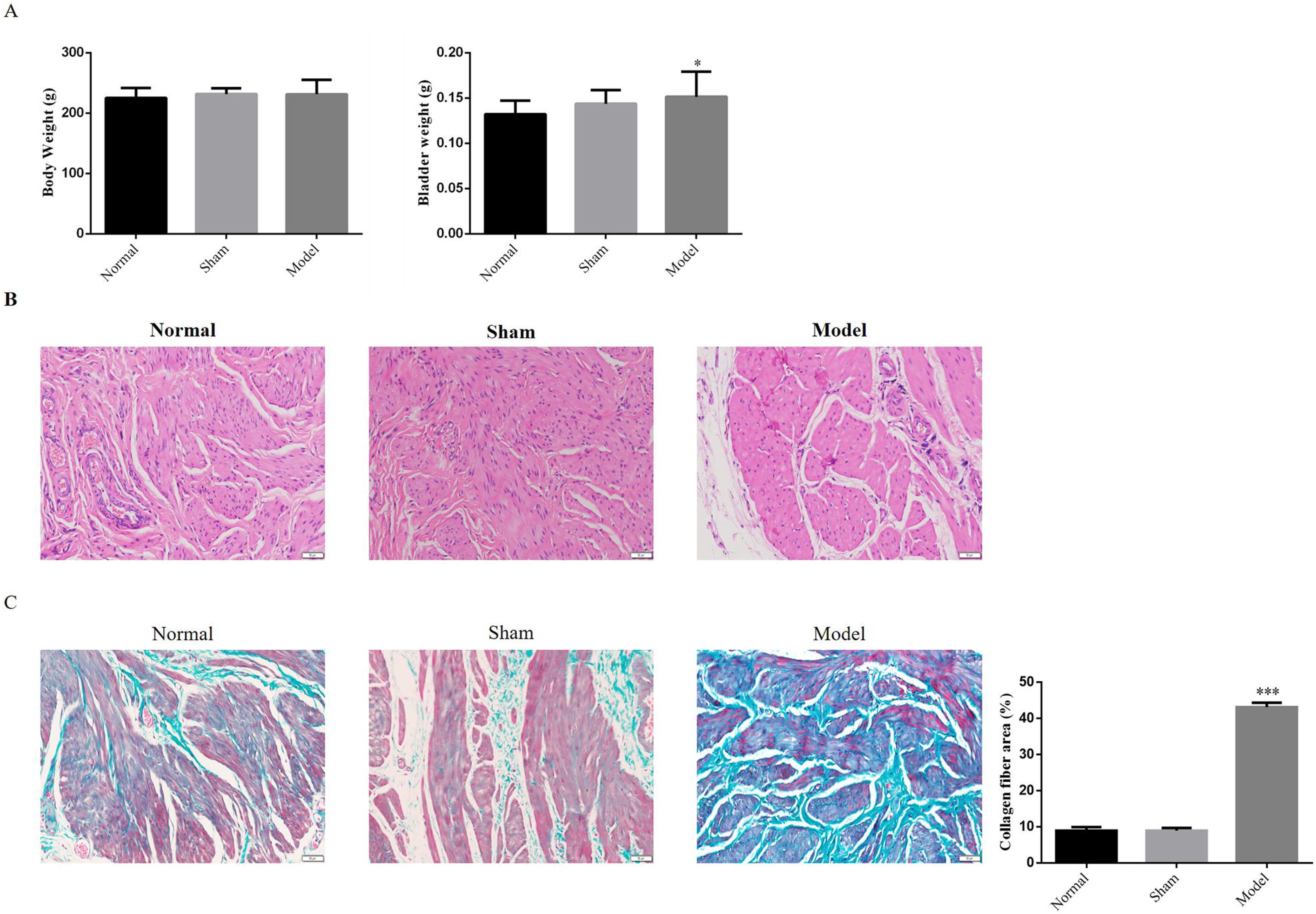
Body weight, bladder weight and pathological changes of rats in different groups A. Body and Bladder weight of difference rats B. The Pathology by HE staining (200×) C. Collagen fibro are of difference rats groups by Masson staining (200×) *: P<0.05, ***: P<0.001, compared with Normal group

### Bladder detrusor cell apoptosis and proliferation of rats in different groups

As revealed by TUNEL staining findings, when compared with Normal group, Sham group was evidently different when it came to the number of apoptotic cells, whereas Model group displayed lower number of such cells (P<0.001, Fig. 2A). After p-histone H3 protein in detrusor tissues was stained, it was found Sham group was not different from Normal group in cell viability, but Model group presented higher cell viability (P<0.001, Fig. 2B). It can be speculated that fibrosis at bladder-urinary tract intersection may be caused by the excessive proliferation when bladder detrusor cell has lost the apoptotic ability.

**Figure 2.**
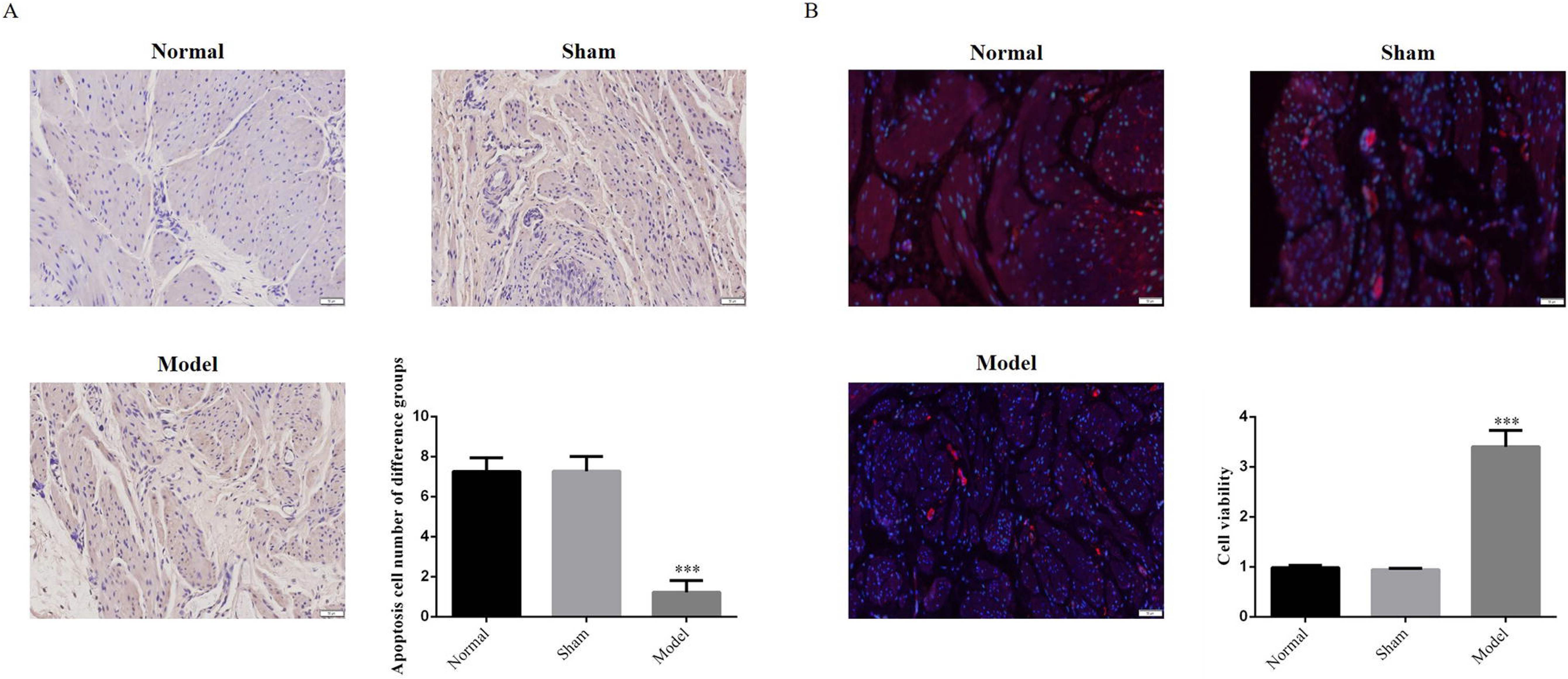
Bladder detrusor cell apoptosis and proliferation of rats in different groups A. Apoptosis cell number of difference groups by TUNEL assay (200×) B. Cell viability of difference groups (200×) ***: P<0.001, compared with Normal group

### ILK protein expression different in bladder tissues of rats

Both WB and IHC findings reveal compared with Normal group, Sham group did not appear quite different in ILK protein expression, but Model group presented significantly higher ILK protein expression level (P<0.001, respectively, Fig. 3A and 3B). It implies fibrosis at urinary tract outlet may remain closely related to the overexpression of ILK.

**Figure 3.**
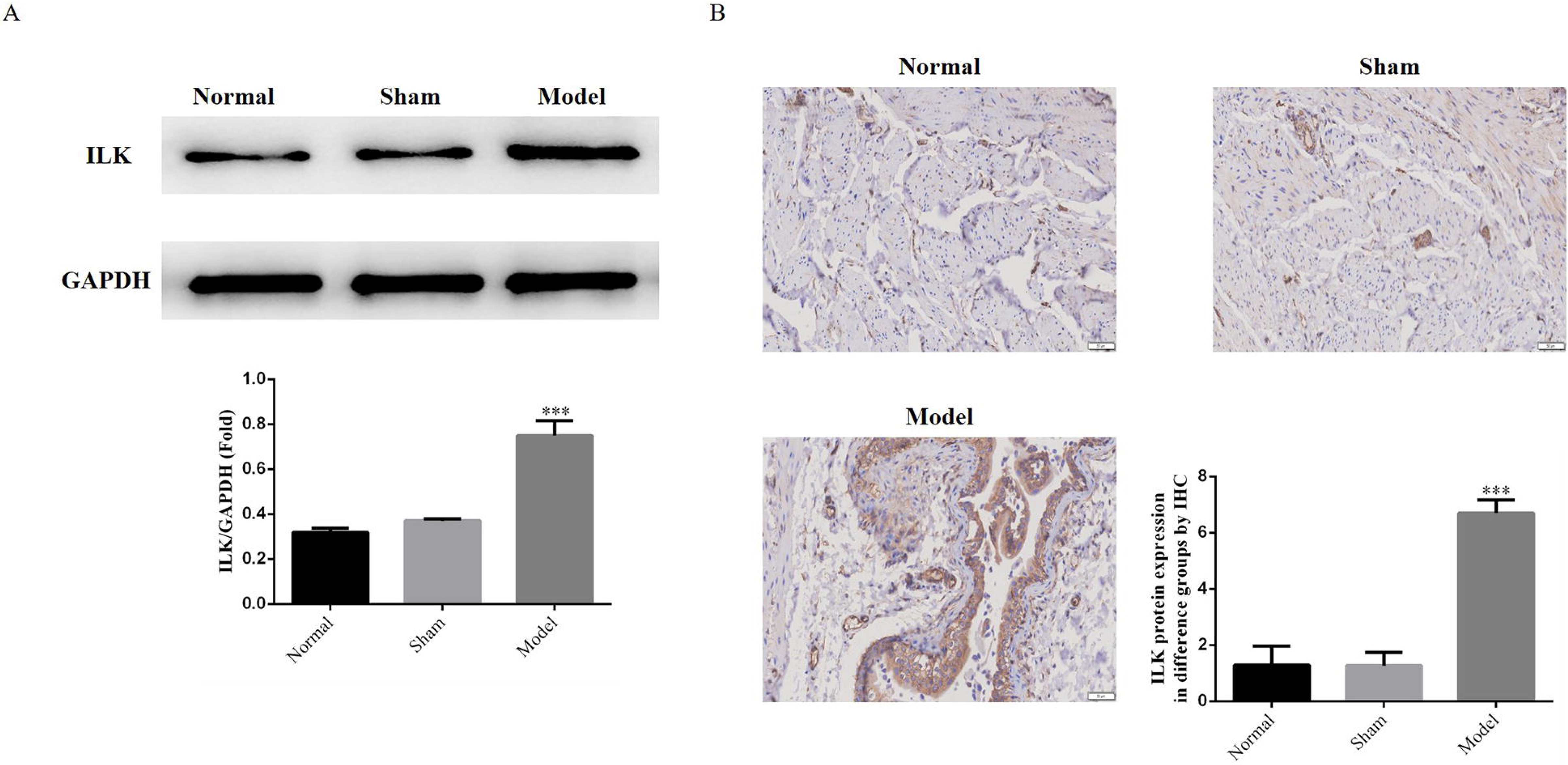
ILK protein expression different in bladder tissues of rats A. ILK protein expression by WB assay B. ILK protein expression by IHC assay in tissues (200×) ***: P<0.001, compared with Normal group

### Primary BSMC cell identification and PBOO cell model construction

α-SMA is a marker protein of BSMC cell. After BSMC cell was extracted, cell immunofluorescence assay reveals both cellular morphology and α-SMA expression are consistent with BSMC criteria (Fig. 4A). TGF-β1 was used to stimulate BSMC and construct a PBOO in-vitro cell model. When stimulated with TGF-β1 at different concentrations, cell proliferation rate and F-actin expression level were greatly elevated when compared with TGF-β1 (0ng/ml) group (P<0.001, Fig. 4B and 4C) and dose-effect relation was identified with TGF-β1 concentration (P<0.05, respectively, Figure 4B and 4C).

**Figure 4.**
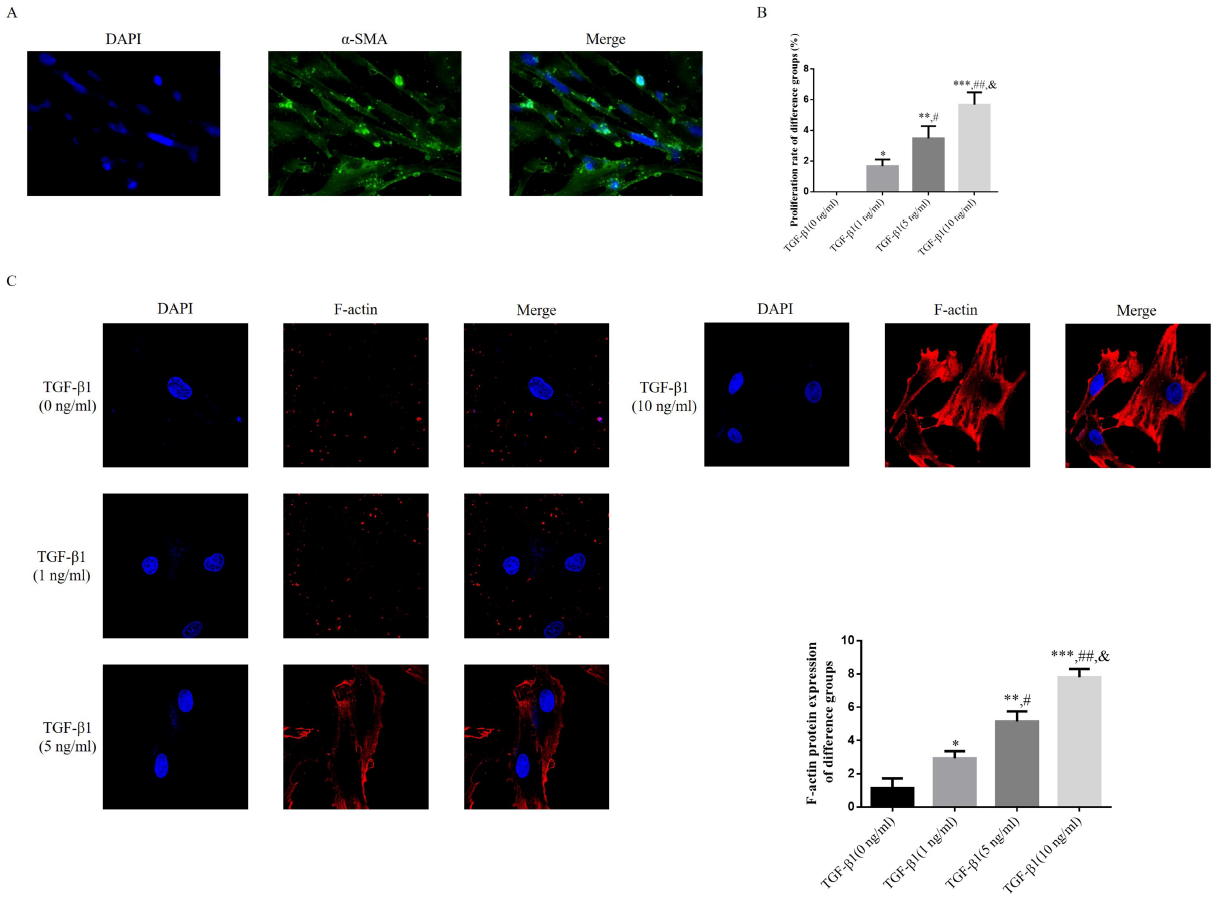
Primary BSMC cell identification and PBOO cell model construction A. α-SMA protein by immunofluorescence assay (200×) B. Proliferation rate of difference groups C. F-actin protein expression of difference groups by immunofluorescence assay (200×) *: P<0.05, **: P<0.01, ***: P<0.001, compared with TGF-β1(0 ng/ml); #: P<0.05, ##: P<0.01, compared with TGF-β1(1 ng/ml; &: P<0.05, compared with TGF-β1(5 ng/ml)

### Expression levels of α-SM-actin and SM22α upon TGF-β1 at various concentrations

The cell immunofluorescence assay indicates compared with TGF-β1 (0mg/ml) group, groups intervened with TGF-β1 showed significantly higher expression levels of α-SM-actin and SM22α (P<0.05, Fig. 5A and 5B) and significant dose-effect relation with TGF-β1 (P<0.05, respectively, Fig. 5A and 5B).

**Figure 5.**
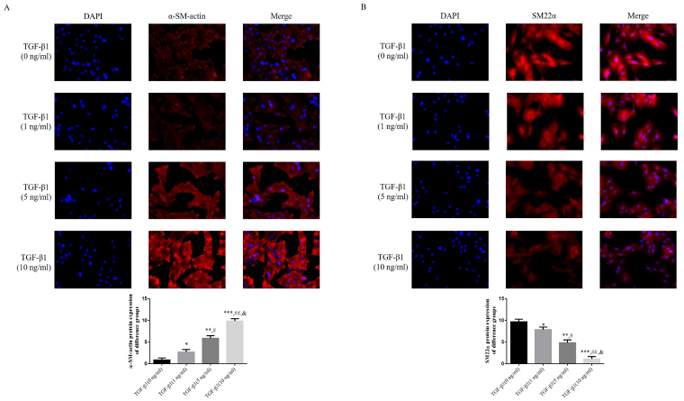
Expression levels of α-SM-actin and SM22α upon TGF-β1 at various concentrations A. α-SM-actin protein expression of difference groups (200×) B. SM-22αprotein expression of difference groups (200×) *: P<0.05, **: P<0.01, ***: P<0.001, compared with TGF-β1(0 ng/ml); #: P<0.05, ##: P<0.01, compared with TGF-β1(1 ng/ml; &: P<0.05, compared with TGF-β1(5 ng/ml)

### Expression of ILK, TLR4, MyD88 and NF-κB(p65) proteins detected with cell immunofluorescence assay

Compared with TGF-β1 (0ng.ml) group, TGF-β1 intervention did raise the expression levels of ILK, TLR4, MyD88 and NF-κB(p65) (P<0.05, respectively, Fig. 6A-6D) and dose-effect relationship with TGF-β1 concentration was detected (P<0.05, respectively, Fig. 6A-6D).

**Figure 6.**
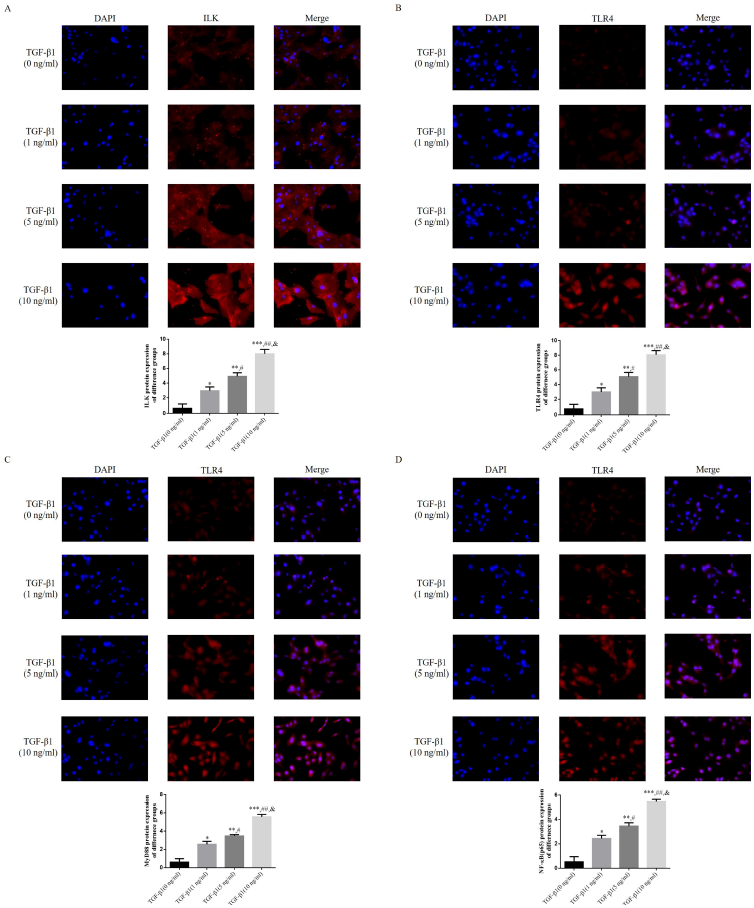
Expression of ILK, TLR4, MyD88 and NF-κB(p65) proteins detected with cell immunofluorescence assay A. ILK protein expression of difference groups (200×) B. TLR4 protein expression of difference groups (200×) C. MyD88 protein expression of difference groups (200×) D. NF-κB(p65) protein expression of difference groups (200×) *: P<0.05, **: P<0.01, ***: P<0.001, compared with TGF-β1(0 ng/ml); #: P<0.05, ##: P<0.01, compared with TGF-β1(1 ng/ml; &: P<0.05, compared with TGF-β1(5 ng/ml)

### Expression of Myocardin and SRF proteins as detected with cell immunofluorescence

Compared with TGF-β1 (0ng.ml) group, TGF-β1 intervention did raise the expression levels of Myocardin and SRF proteins (P<0.05, respectively, Fig. 7A and 7B) and dose-effect relationship with TGF-β1 concentration was detected (P<0.05, respectively, Fig. 7A and 7B).

**Figure 7.**
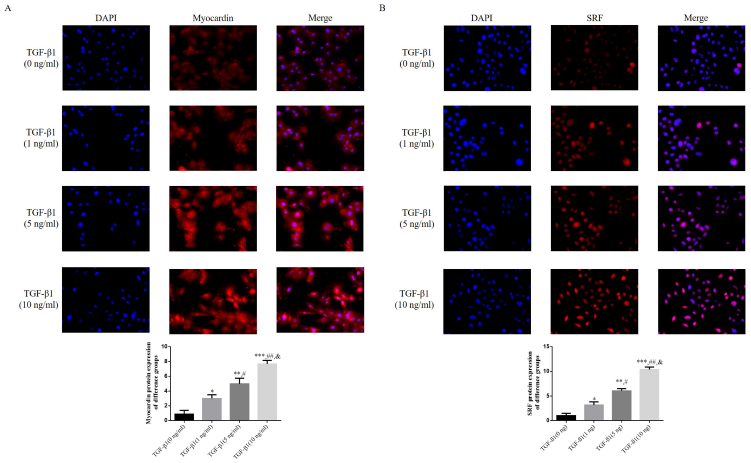
Expression of Myocardin and SRF proteins as detected with cell immunofluorescence A. Myocardin protein expression of difference groups (200×) B. SRF protein expression of difference groups (200×) *: P<0.05, **: P<0.01, ***: P<0.001, compared with TGF-β1(0 ng/ml); #: P<0.05, ##: P<0.01, compared with TGF-β1(1 ng/ml; &: P<0.05, compared with TGF-β1(5 ng/ml)

### Differences in the expression of ILK, TLR4, MyD88, NF-κB(p65), Myocardin and SRF

Compared with TGF-β1(0ng.ml) group, TGF-β1 intervention did raise the expression levels of ILK, TLR4, MyD88, NF-κB(p65), Myocardin and SRF (P<0.05, respectively, Fig. 8A and 8B) and dose-effect relationship with TGF-β1 concentration was detected (P<0.05, respectively, Fig. 8A and 8B).

**Figure 8.**
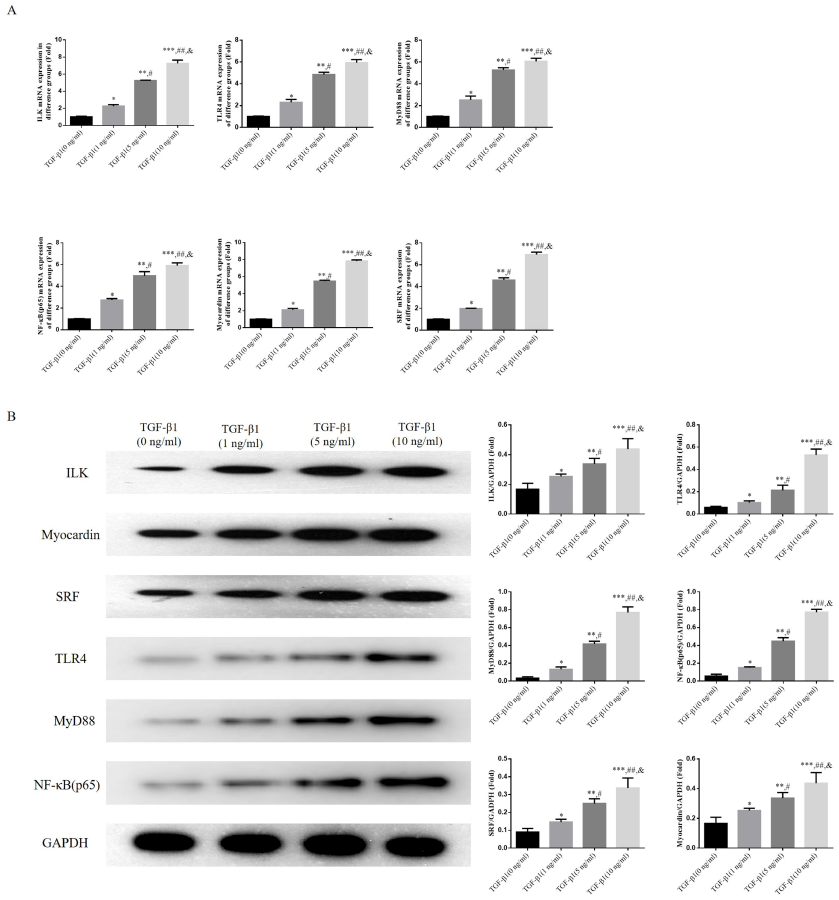
Differences in the expression of ILK, TLR4, MyD88, NF-κB(p65), Myocardin and SRF A. Relative mRNA expression in difference groups B. Relative proteins expression in difference groups *: P<0.05, **: P<0.01, ***: P<0.001, compared with TGF-β1(0 ng/ml); #: P<0.05, ##: P<0.01, compared with TGF-β1(1 ng/ml; &: P<0.05, compared with TGF-β1(5 ng/ml)

### Role of ILK in BSMC proliferation and F-action expression

When compared with treatment-free BDMC group, LV-ILK group, TGF-β1 group, TGF-β1+LV-NC group and TGF-β1+LV-ILK group showed significantly higher proliferation rates (P<0.01, respectively, Fig. 9A) and higher F-actin expression levels (P<0.01, respectively, Fig. 8B). And when compared with TGF-β1 group, TGF-β1+LV-ILK group showed significantly higher proliferation rate (P<0.05, respectively, Fig. 9A) and F-actin expression level as well (P<0.05, respectively, Fig. 9B).

**Figure 9.**
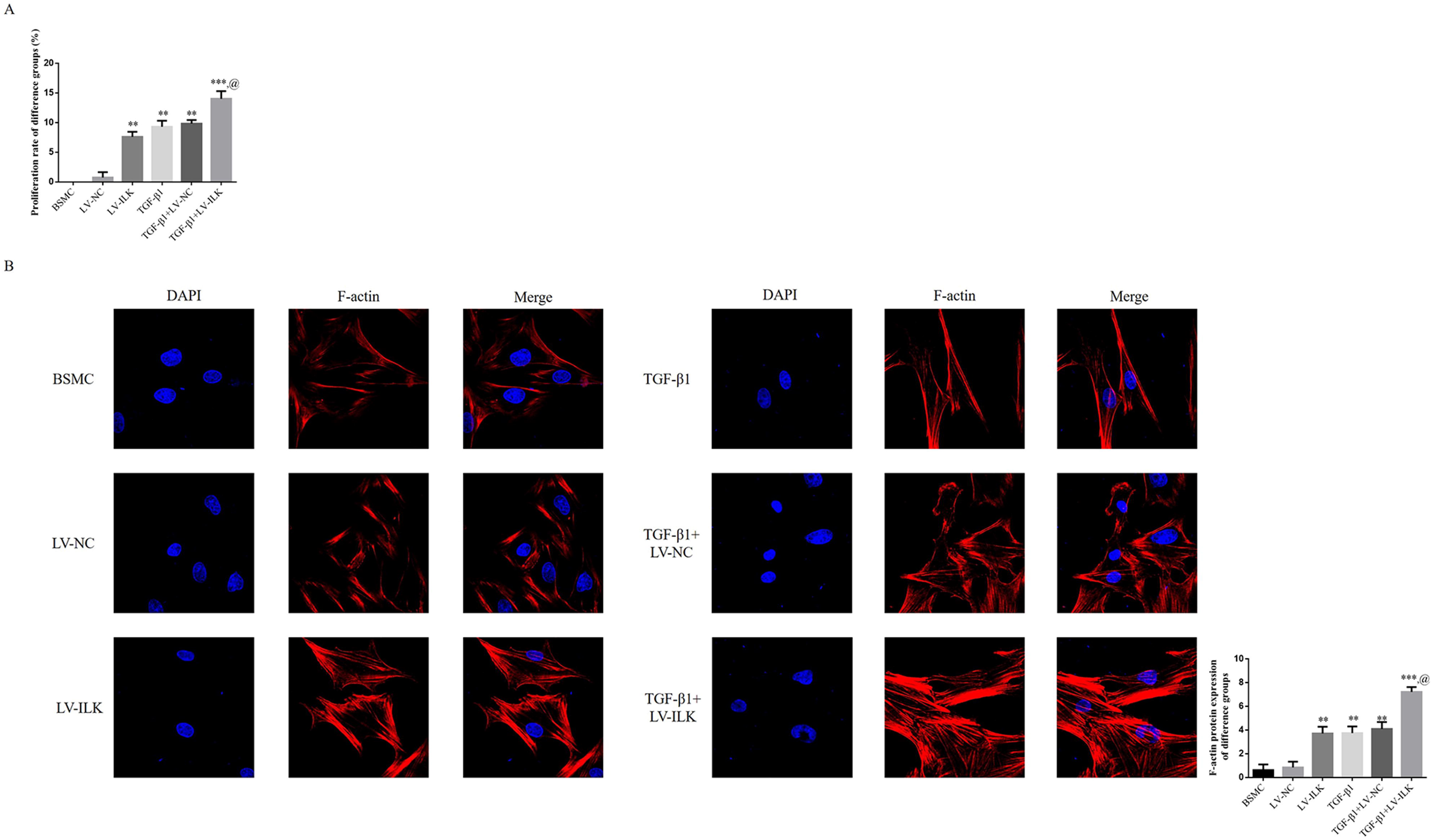
Role of ILK in BSMC proliferation and F-action expression A. Proliferation rate of difference groups B. F-actin protein expression of difference groups (200×) **: P<0.01; ***: P<0.001, compared with BSMC group; @: P<0.05, compared with TGF-β1 group

### Effect of ILK onα-SM-actin and SM22α

When compared with treatment-free BDMC group, LV-ILK group, TGF-β1 group, TGF-β1+LV-NC group and TGF-β1+LV-ILK group showed significantly higher α-SM-actin expression levels (P<0.01, respectively, Fig. 10A) but lower SM22α expression levels (P<0.01, respectively, Fig. 10B). And when compared with TGF-β1 group, TGF-β1+LV-ILK group showed significantly higher α-SM-actin expression level (P<0.05, respectively, Fig. 10A) but lower SM22α protein expression level (P<0.05, respectively, Fig. 10B).

**Figure 10.**
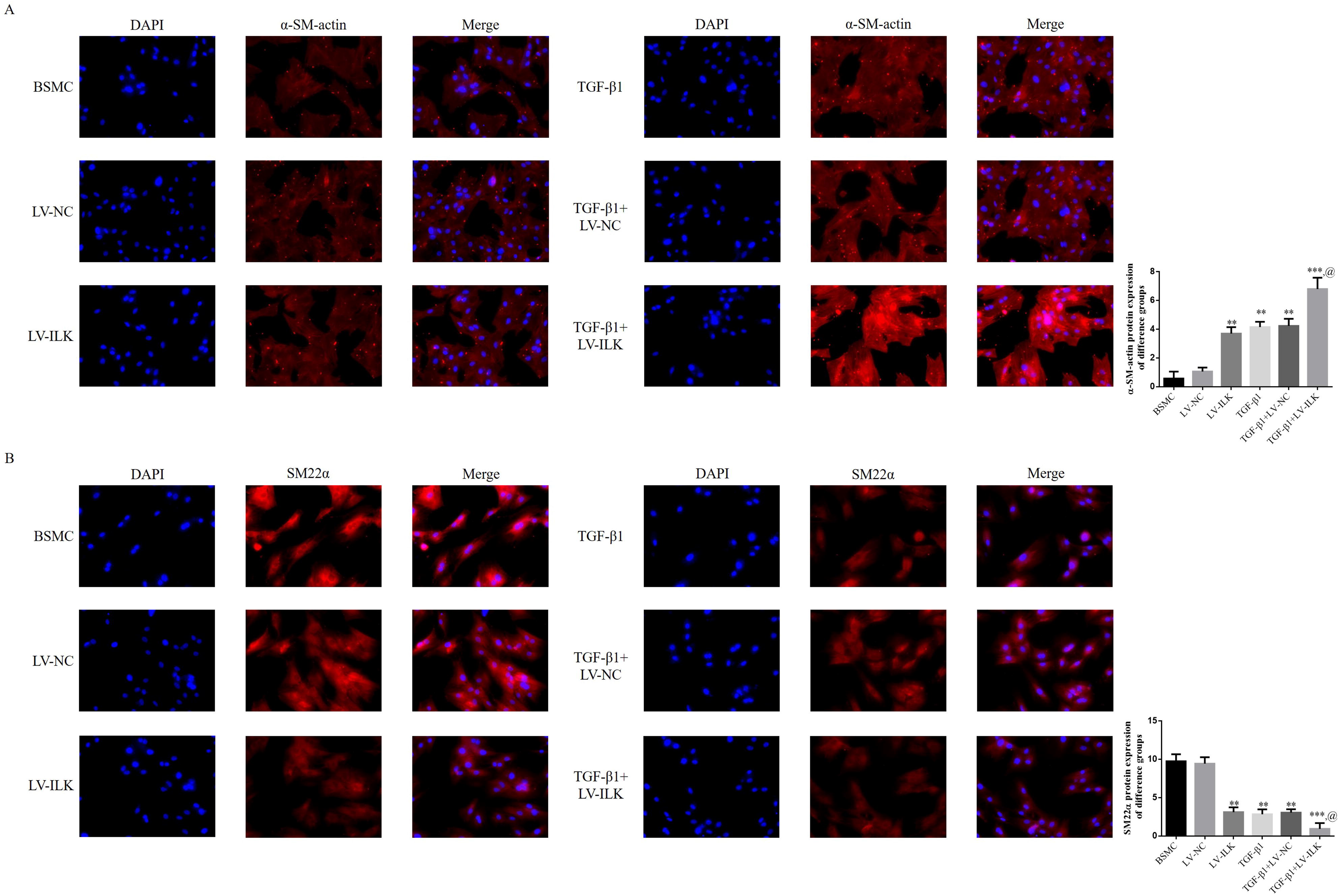
Effect of ILK on α-SM-actin and SM22α A. α-SMA protein expression of difference groups (200×) B. SM22α protein expression of difference groups (200×) **: P<0.01; ***: P<0.001, compared with BSMC group; @: P<0.05, compared with TGF-β1 group

### Expression of ILK, TLR4, MyD88 and NF-κB(p65) proteins

When compared with treatment-free BDMC group, LV-ILK group, TGF-β1 group, TGF-β1+LV-NC group and TGF-β1+LV-ILK group showed significantly higher expression levels of ILK, TLR4, MyD88 and NF-κB(p65) proteins (P<0.01, respectively, Fig. 11A-11D). And when compared with TGF-β1 group, TGF-β1+LV-ILK group showed significantly higher expression levels of ILK, TLR4, MyD88 and NF-κB(p65) proteins (P<0.05, respectively, Fig. 11A-11D).

**Figure 11.**
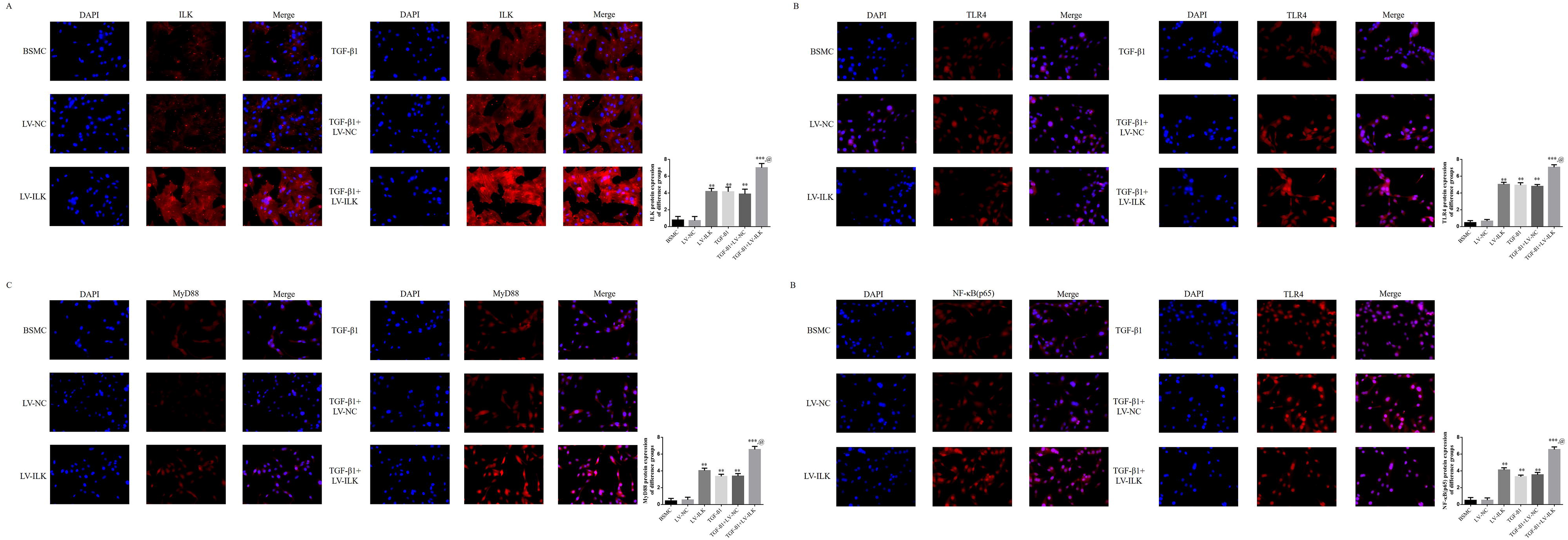
Expression of ILK, TLR4, MyD88 and NF-κB(p65) proteins A. ILK protein expression of difference groups (200×) B. TLR4 protein expression of difference groups (200×) C. MyD88 protein expression of difference groups (200×) D. NF-κB(p65) protein expression of difference groups (200×) **: P<0.01; ***: P<0.001, compared with BSMC group; @: P<0.05, compared with TGF-β1 group

### Effect of ILK overexpression on Myocardin and SRF proteins’ expression

When compared with treatment-free BDMC group, LV-ILK group, TGF-β1 group, TGF-β1+LV-NC group and TGF-β1+LV-ILK group showed significantly higher expression levels of Myocardin and SRF proteins (P<0.01, respectively, Fig. 12A and 12B). And when compared with TGF-β1 group, TGF-β1+LV-ILK group showed significantly higher expression levels of Myocardin and SRF proteins (P<0.05, respectively, Fig. 12A and 12B).

**Figure 12.**
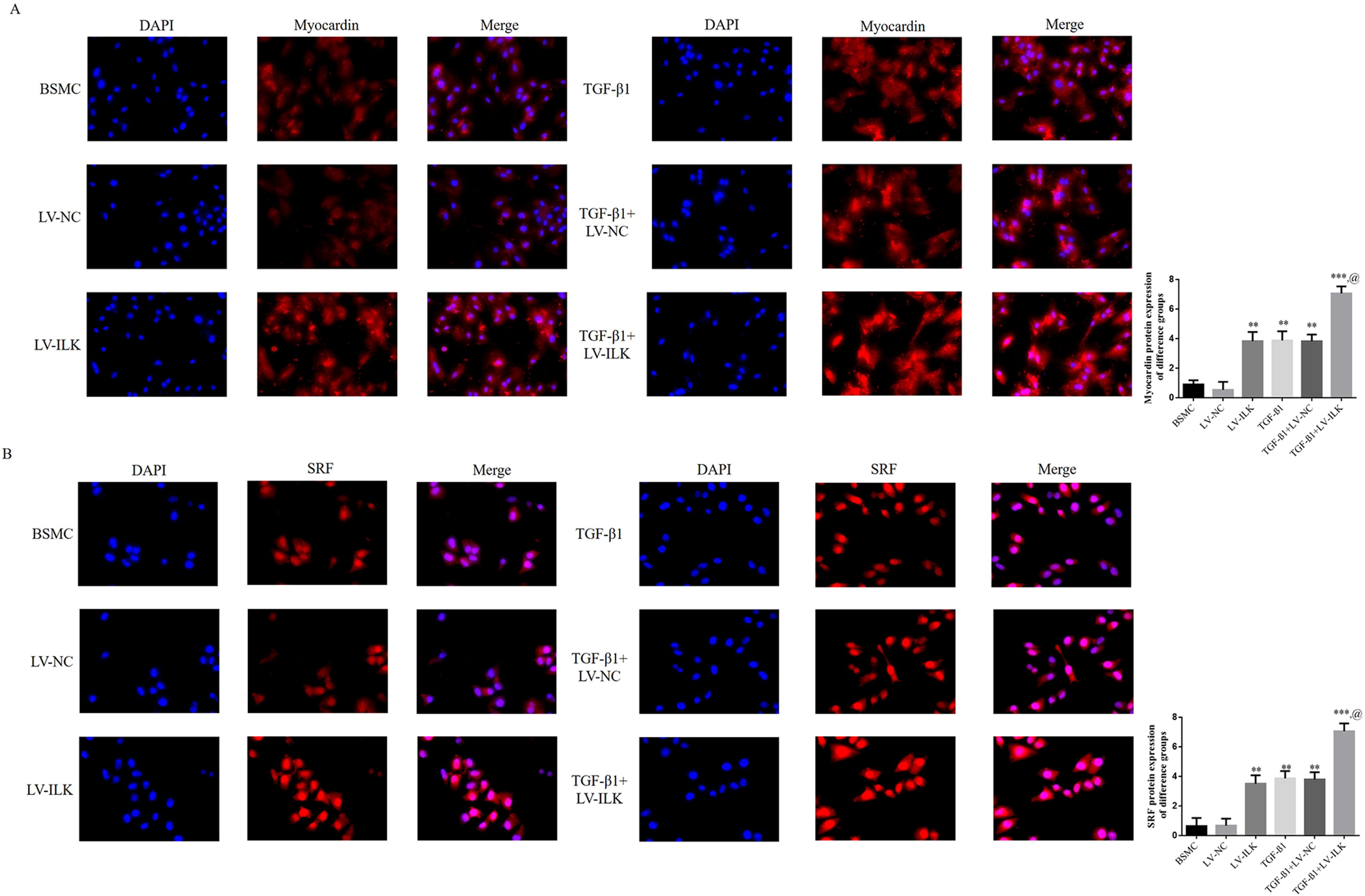
Effect of ILK overexpression on Myocardin and SRF proteins’ expression A. Myocardin protein expression of difference groups (200×) B. SRF protein expression of difference groups (200×) **: P<0.01; ***: P<0.001, compared with BSMC group; @: P<0.05, compared with TGF-β1 group

### Effect of ILK overexpression on related genes and proteins

As revealed by RT-PCR and WB assays, compared with treatment-free BDMC group, LV-ILK group, TGF-β1 group, TGF-β1+LV-NC group and TGF-β1+LV-ILK group showed significantly higher expression levels of ILK, TLR4, MyD88, NF-κB(p65), Myocardin and SRF (P<0.01, respectively, Fig. 13A and 13B). And when compared with TGF-β1 group, TGF-β1+LV-ILK group showed significantly higher expression levels of ILK, TLR4, MyD88, NF-κB(p65), Myocardin and SRF genes and proteins (P<0.05, respectively, Fig. 13A and 13B).

**Figure 13.**
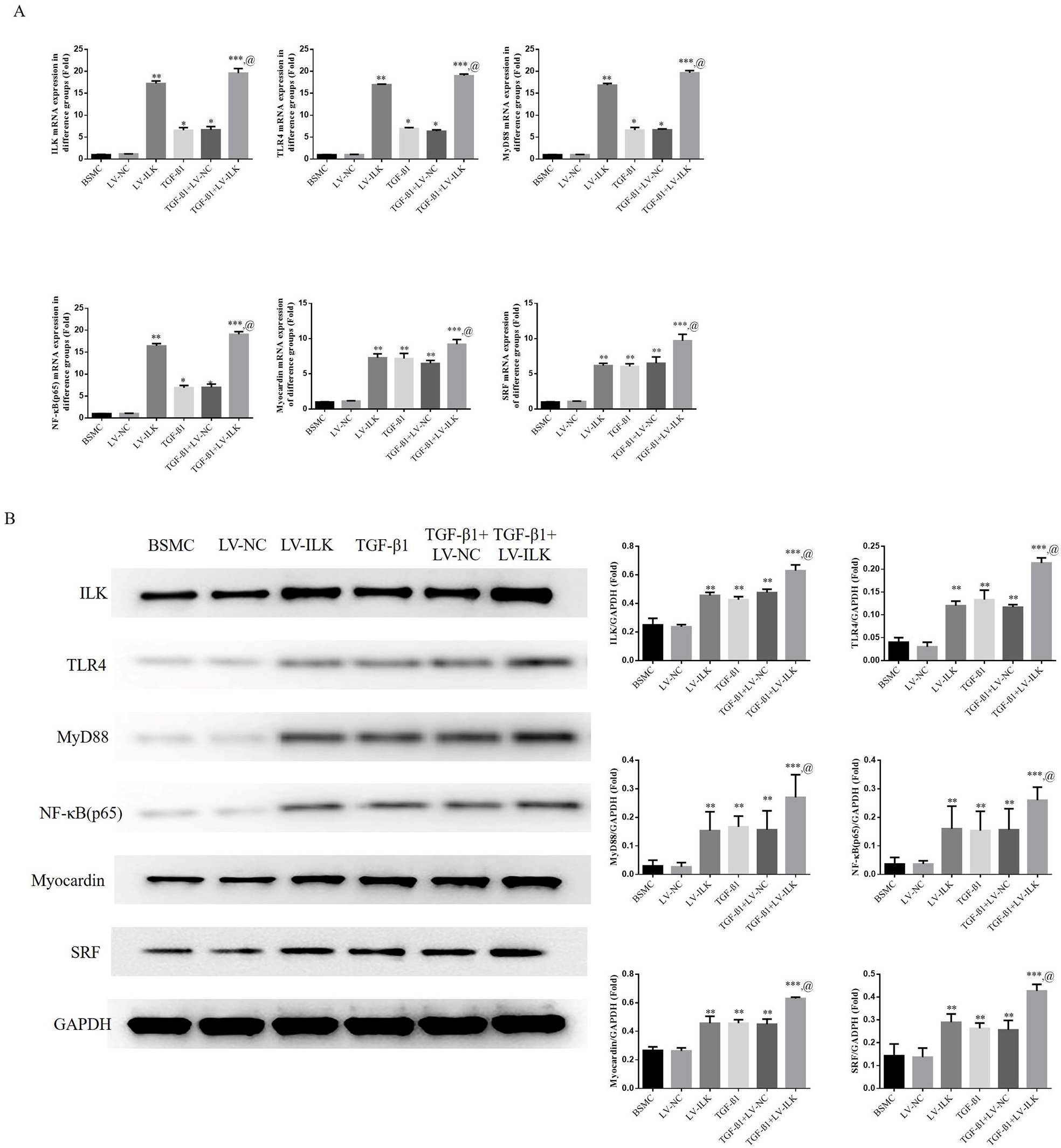
Effect of ILK overexpression on related genes and proteins A. Relative mRNA expression in difference groups B. Relative protein expression in difference groups **: P<0.01; ***: P<0.001, compared with BSMC group; @: P<0.05, compared with TGF-β1 group

### Effect of ILK knockout on cell proliferation and F-action expression

In order to verify the effect of si-ILK, RT-PCR assay was performed. The results indicate when compared with treatment-free BDMC group, si-ILK-1, si-ILK-2 and si-ILK-3 groups displayed significantly lower ILK expression levels (P<0.05, respectively, Fig. 14A) with si-ILK-2 yielding the best inhibiting effect. When compared with treatment-free BDMC group, si-ILK group had significantly lower cell proliferation rate (P<0.001, Fig. 14B), whereas TGF-β1 and TGF-β1+si-NC groups displayed higher cell proliferation rates (P<0.001, Fig. 14B). As suggested by cell immunofluorescence assay, when compared with treatment-free BDMC group, si-ILK group had significantly lower F-actin expression level (P<0.001, Fig. 14C), but TGF-β1 group and TGF-β1+si-NC group showed higher F-actin expression levels (P<0.001, respectively, Fig. 14C). When compared with TGF-β1 group, TGF-β1+si-ILK group had lower F-actin expression level (P<0.001, Fig. 14C).

**Figure 14.**
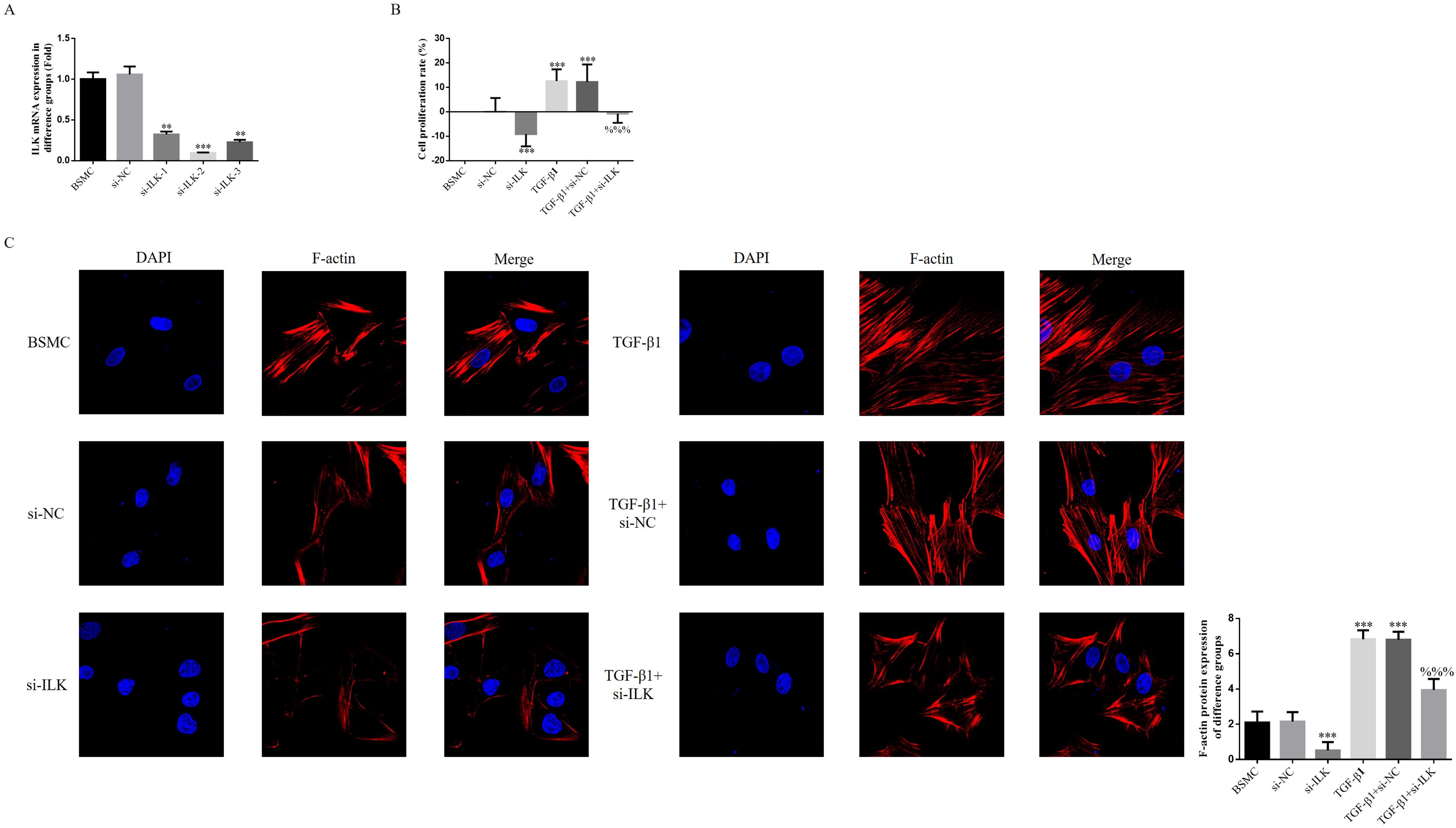
Effect of ILK knockout on cell proliferation and F-action expression A. ILK mRNA expression in difference groups (Fold) B. Cell proliferation by CCK-8 assay C. F-actin protein expression of difference groups (200×) ***: P<0.001, compared with BSMC group; %%%: P<0.001, compared with TGF-β1 group

### Effect of ILK knockout on expression ofα-SM-actin and SM22α proteins

When compared with BSMC group, si-ILK group displayed lower α-SM-actin expression but higher SM22α expression (P<0.001, respectively, Fig. 15A and 15B), whereas TGF-β1 group and TGF-β1+si-NC group presented higher α-SM-actin expression but lower SM22α expression (P<0.001, respectively, Fig. 15A and 15B). And when compared with TGF-β1 group, TGF-β1+si-ILK presented lower α-SM-actin expression but higher SM22α expression (P<0.001, respectively, Fig. 15A and 15B).

**Figure 15.**
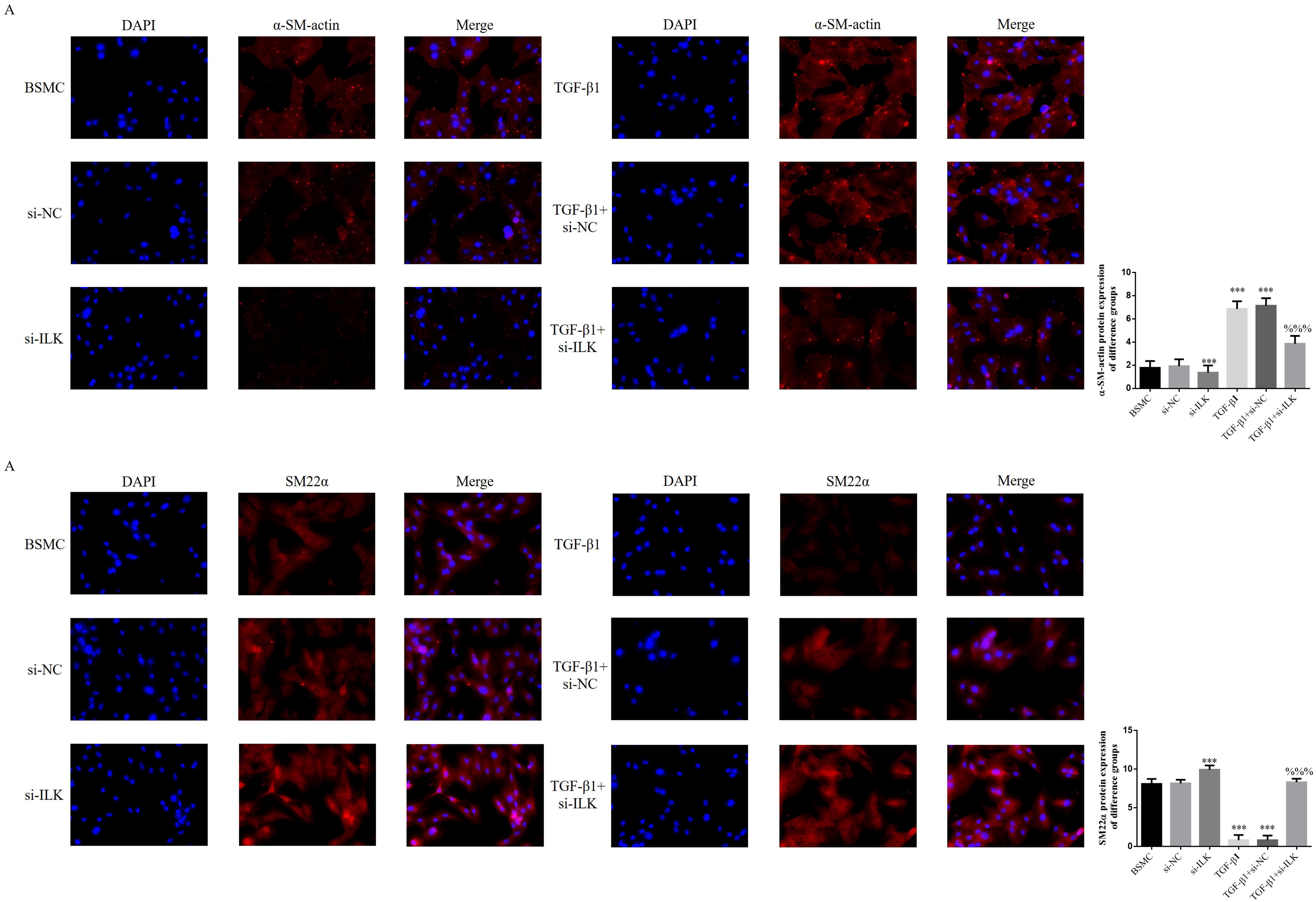
Effect of ILK knockout on expression of α-SM-actin and SM22α proteins A. α-SM-actin protein expression of difference groups (200×) B. SM-22α protein expression of difference groups (200×) ***: P<0.001, compared with BSMC group; %%%: P<0.001, compared with TGF-β1 group

### Effect of ILK knockout on related proteins

When compared with BSMC group, si-ILK group displayed lower expression levels of ILK, TLR4, MyD88, NF-κB(p65), Myocardin and SRF (P<0.001, respectively, Fig. 17A and 17B), whereas TGF-β1 group and TGF-β1+si-NC group presented higher expression levels of ILK, TLR4, MyD88, NF-κB(p65), Myocardin and SRF (P<0.001, respectively, Fig. 16A-16D). And when compared with TGF-β1 group, TGF-β1+si-ILK presented lower expression levels of ILK, TLR4, MyD88, NF-κB(p65), Myocardin and SRF (P<0.001, respectively, Fig. 16A-16D, Fig. 17A and 17B).

**Figure 16.**
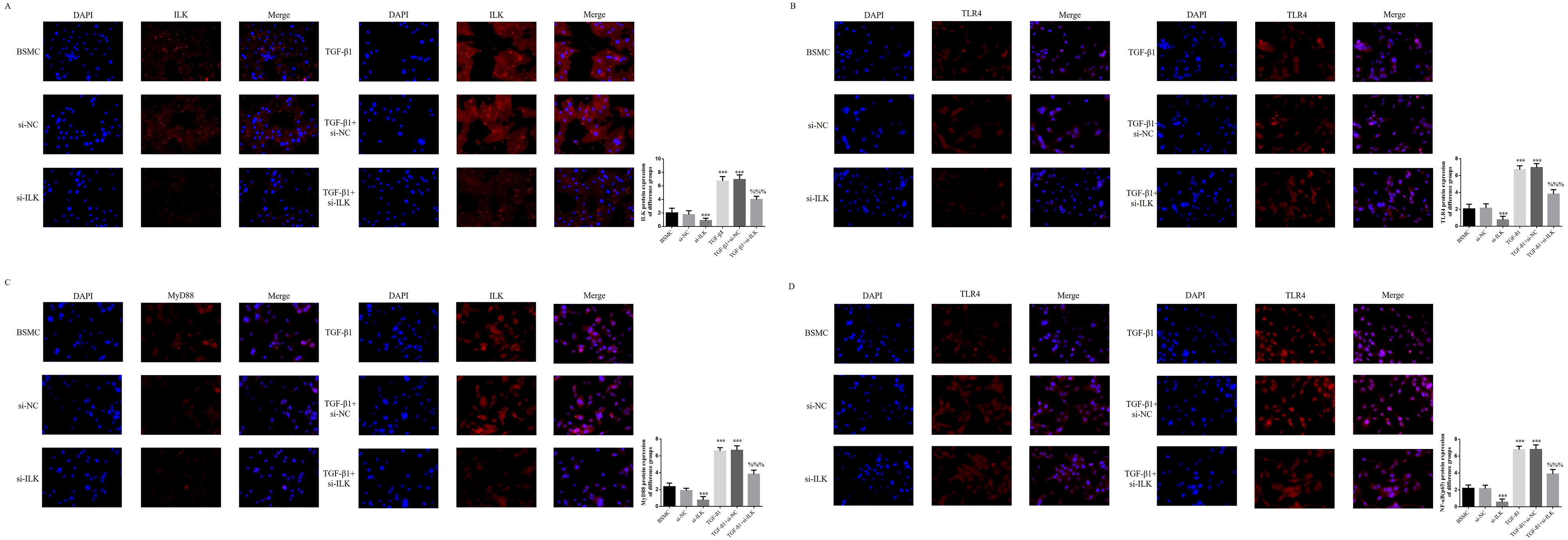
Effect of ILK knockout on related proteins A. ILK protein expression of difference groups (200×) B. TLR4 protein expression of difference groups (200×) C. MyD88 protein expression of difference groups (200×) D. NF-κB(p65) protein expression of difference groups (200×) ***: P<0.001, compared with BSMC group; %%%: P<0.001, compared with TGF-β1 group

**Figure 17.**
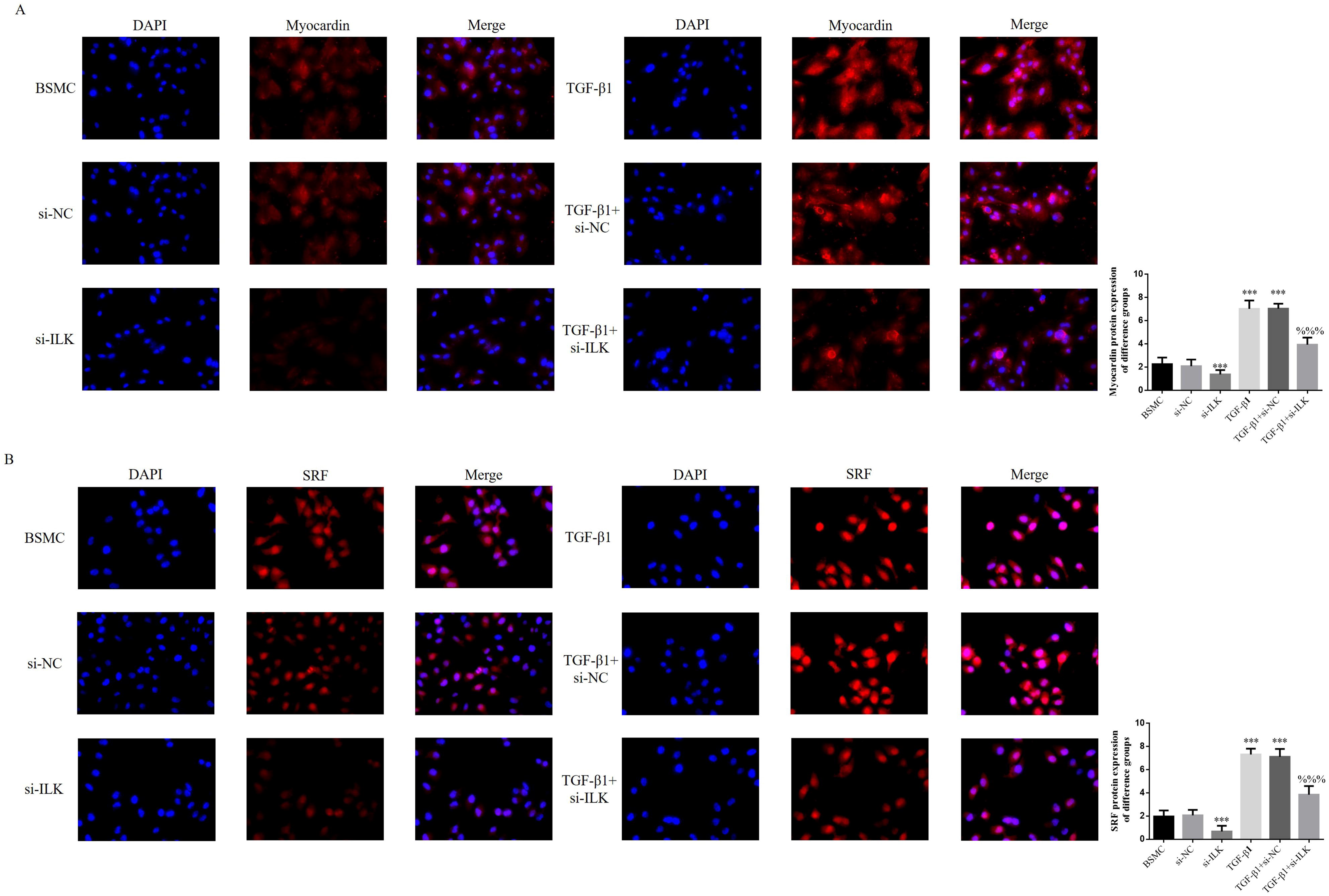
Myocardin and SRF protein expression A. Myocardin protein expression of difference groups (200×) B. SRF protein expression of difference groups (200×) ***: P<0.001, compared with BSMC group; %%%: P<0.001, compared with TGF-β1 group

### Effect of ILK knockout on expression of related genes and proteins

When compared with BSMC group, si-ILK group displayed lower expression levels of ILK, TLR4, MyD88, NF-κB(p65), Myocardin and SRF genes and proteins (P<0.001, respectively, Fig. 18A and 18B), whereas TGF-β1 group and TGF-β1+si-NC group presented higher expression levels of ILK, TLR4, MyD88, NF-κB(p65), Myocardin and SRF genes and proteins (P<0.001, respectively, Fig. 18A and 18B). And when compared with TGF-β1 group, TGF-β1+si-ILK presented lower expression levels of ILK, TLR4, MyD88, NF-κB(p65), Myocardin and SRF genes and proteins (P<0.001, respectively, Fig. 18A and 18B).

**Figure 18.**
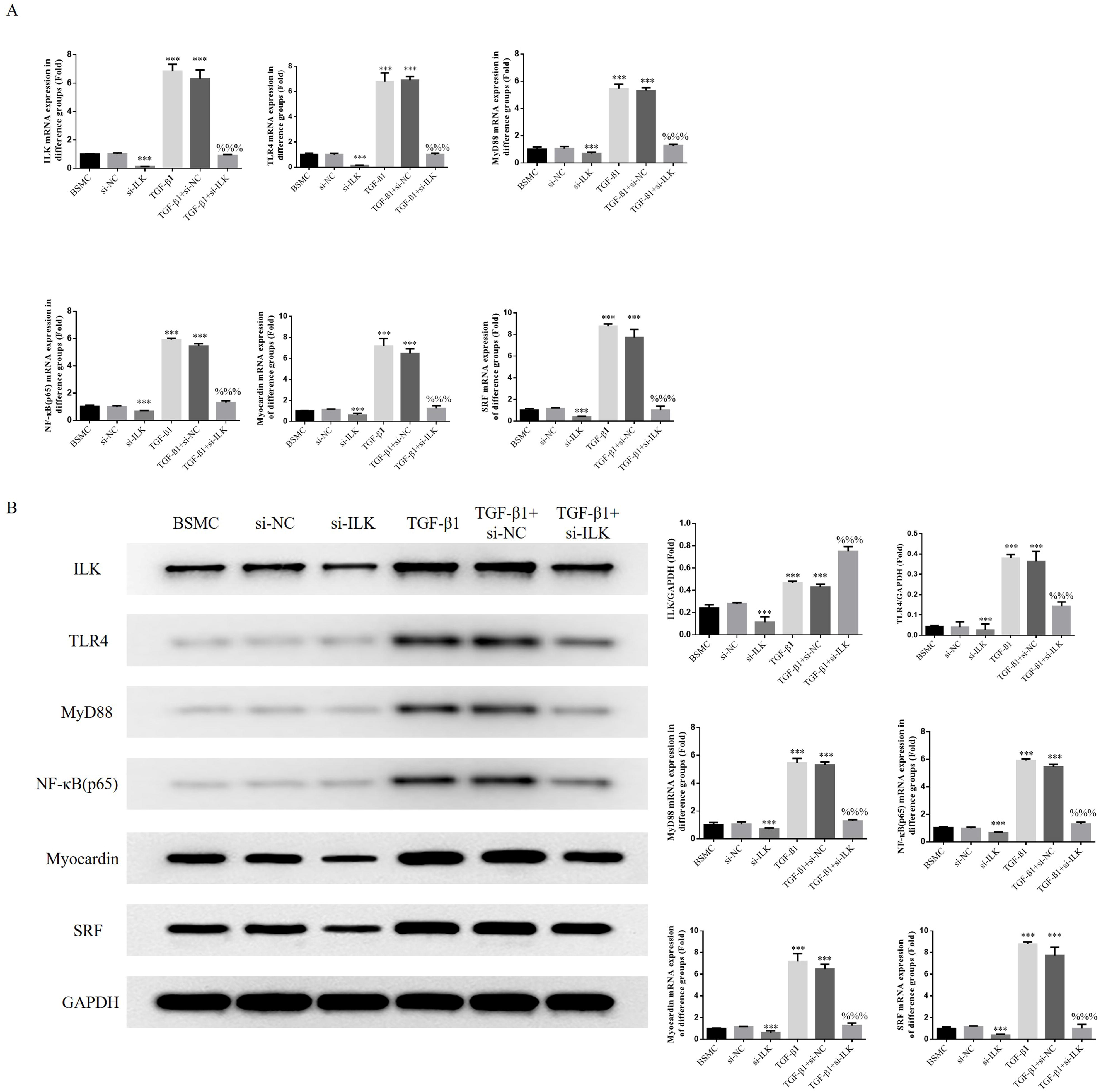
Relative mRNA and protein expression A. Relative mRNA expression B. Relative protein expression ***: P<0.001, compared with BSMC group; %%%: P<0.001, compared with TGF-β1 group

## Discussion

PBOO is a generic term to the diseases concerning dysuria arising from elevated urine outflow tract resistance when bladder and/or urethra are affected by various factors [8]. When examining the pathological changes and fibrosis of constructed PBOO rat model, the present study finds out when cell apoptosis-proliferation balance of smooth muscle at bladder outlet of PBOO rats gets undermined, fibrosis rises sharply. The finding suggest excessive proliferation of smooth muscle may be an important factor leading to PBOO occurrence.

Researches indicate harmful stimuli may induce BSMC to undergo a string of adaptive changes including cell hypertrophy, phenotypic transformation, proliferation and apoptosis, and finally the functional decompensation of bladder. Upon an obstruction, BSMC may experience reversible phenotypic transformation and changes in related genes once being damaged or stimulated so that the cell contraction protein is less expressed and BSMC proliferate more to impair the contraction function [9]. The vasoactive substances generated by local inflammatory cells of bladder after obstruction occurs, including transforming growth factor-β1 (TGF-β1), bFGF, and HB-EGF, could promote BSMC proliferation [10]. Many scholars including Luo have testified BSMC does undergo phenotypic transformation and then changes its cell proliferation state and secrete extracellular matrix (ECM) to lead to bladder fibrosis [11]. Thus, obstruction-induced BSMC phenotypic serves as an initial link in the proliferative change and migration of BSMC. In present study, we stimulate primary BSMC with TGF-β1 to construct a in-vitro PBOO cell model. Our findings suggest both PBOO proliferation rate and fibrosis are evidently elevated.

Integrin-linked kinase (ILK) is a protein with molecular weight being 59kD discovered by Hannigan in 1996 when he studied β1 integrin-binding protein. It is a serine/threonine protein kinase that functions to mediate the signal pathway function within integrin cell, being expressed in many cells of mammals [12]. ILK binds with the subunit cytoplasm of integrin β. The extracellular region of integrin connected with extracellular matrix plays a part in several physiological functions. Wu et al demonstrated through their study that ILK could negatively regulate RhoA/ROCK activity and become involved in expression of SMC phenotypic transformation marker gene. Besides, it also plays a vital part in maintaining SMC contraction phenotype [13–15]. In our study, we extracted primary BSMC from rats’ bladder outflow tissue, and then employed TGF-β1 to stimulate and construct PBOO cell model. The results indicate after being stimulated by TGF-β1, BSMC exhibited the pathological changes similar to that of animal model. Transfection of ILK into the cell brought in excessive proliferation and higher fibrosis rate of BSMC. However, when ILK got knocked out, the excessive proliferation and fibrosis of BSMC induced by TGF-β1 were significantly improved. It can be inferred that PBOO occurrence is closely related to the overexpression of ILK.

TLR4/MyD88/NF-κB(p65) signal pathway activation has great significance in inflammation and fibrosis processes [16–18]. Some studies reveal ILK could effectively regulate this pathway [19]. On the other hand, it has been discovered by recent studies that TLR4/MyD88/NF-κB(p65) signal pathway could effectively regulate expression of SRF and Myocardin [20]. Olson Lab detected the most effective SRF coactivator by far-Myocardin which could initialize the phenotypic transformation procedure of smooth muscle cell (SMC) and activate the transcription of a series of SRF-regulated and codified cell scaffolding proteins and contraction-associated proteins so as to regulate the SMC phenotyping [20]. In this study, the findings suggest upon ILK overexpression, TGF-β1 stimulation and TLR4/MyD88/NF-κB(p65) activation, downstream SRF and Myocardin gene and proteins change in their expression. Apart from that, BSMC cell proliferation and fibrosis would be promoted also. However, if ILK is knocked out, the excessive proliferation and fibrosis of BSMC induced by high TGF-β1 get inhibited. Along with the reduced activity of TLR4/MyD88/NF-κB(p65), downstream SRF and Myocardin expression are also undermined.

In conclusion, this study employs both in vivo and in vitro experiments to prove the important role played by ILK overexpression in BPOO genesis and development. When ILK gets knocked down, the BPOO-induced excessive proliferation and cell fibrosis are both ameliorated. The action mechanism may stay closely related to the TLR4/MyD88/NF-κB(p65) and downstream SRF and Myocardin expression.

